# Multi-Algorithm Machine Learning Benchmarking for Pan-Cancer Classification from Tumour-Educated Platelet RNA Sequencing

**DOI:** 10.64898/2026.05.22.727079

**Authors:** Devam Himanshu Zalawadia, Vinaya Bhate, Tanesha Deepan Chakravarthy, Aushdhan Govindraj Chetty, Sonalika Ray

## Abstract

Tumour-educated platelets (TEPs) carry cancer-type-specific RNA signatures accessible through whole-blood RNA sequencing, but systematic multi-algorithm benchmarking with quantified statistical uncertainty had not been applied to the GSE68086 dataset, the field’s primary reference cohort. We applied an end-to-end transcriptomic and machine learning framework to 280 whole-blood platelet RNA-seq samples from six cancer types (non-small cell lung cancer, colorectal cancer, glioblastoma multiforme, hepatobiliary cancer, breast cancer, and pancreatic cancer) and healthy donors. After a standardised preprocessing and normalisation pipeline, seven supervised classifiers - Logistic Regression, SVM (RBF), XGBoost, LightGBM, Random Forest, K-Nearest Neighbours, and a Multilayer Perceptron were benchmarked using stratified 5-fold cross-validation and a held-out test set. Statistical uncertainty was quantified via 2,000-resample percentile bootstrap confidence intervals. Multinomial Logistic Regression achieved the highest test macro F1-score (0.522) and macro-averaged ROC-AUC (0.869), both substantially above the seven-class chance level (1/7 ≈ 0.14). SHAP analysis of the Random Forest classifier identified IFITM3 as the globally dominant TEP biomarker; cancer-type-specific discriminators included ATP5PD (hepatobiliary cancer), C6orf62 (NSCLC and pancreatic cancer), VPS13C (healthy donors), and TMSB4Y (breast cancer). Gene Ontology and KEGG pathway enrichment corroborated the biological specificity of identified transcriptomic signatures. These results support the diagnostic potential of TEP transcriptomics as a multi-class liquid biopsy platform and provide a methodologically transparent, reproducible reference framework for future blood-based cancer classification studies.

## 1. Introduction

Cancer remains a leading cause of premature mortality worldwide, 19.97 million new diagnoses and 9.74 million deaths were recorded globally in 2022 alone [1]. Despite advances in targeted therapy and immunotherapy, the prognosis of most solid tumours is dictated primarily by disease stage at diagnosis [2, 1]. The problem is that most cancers surface at locally advanced or metastatic stages, when curative intervention is far more difficult. Early-stage disease is frequently asymptomatic, and current screening modalities lack the sensitivity and specificity needed for population-wide application [3]. Conventional diagnostics, radiological imaging and tissue biopsy - carry procedural risk, substantial cost, and an inherent inability to track tumour heterogeneity across serial time points [4]. These constraints have driven interest in minimally invasive, blood-based approaches that can detect cancer early and, more critically, discriminate among multiple tumour types from a single clinical sample [3, 4].

Liquid biopsy - molecular interrogation of tumour-derived circulating components has transformed cancer diagnostics and disease monitoring [4]. Circulating tumour DNA (ctDNA), circulating tumour cells (CTCs), cell-free RNA, and tumour-derived extracellular vesicles each encode distinct genomic, transcriptomic, and epigenomic information reflecting the molecular state of the primary tumour and its metastases [3, 4]. Unlike tissue biopsy, liquid biopsy is compatible with serial sampling, enabling real-time therapeutic monitoring and early detection of acquired resistance [4]. ctDNA-based assays, however, lose sensitivity in early-stage disease when tumour shedding is low and the tumour fraction in circulating cell-free DNA is minimal [3]. Robust multi-class discrimination, determining not merely that cancer is present but which tumour type remains technically and computationally challenging for most circulating analyte platforms [1, 4]. Blood platelets, classically defined as anucleate mediators of haemostasis, are active participants in tumour biology [5]. During circulation through the tumour-associated vasculature, platelets sequester tumour-derived growth factors, proteases, nucleic acids, and signalling intermediates through dynamic crosstalk with tumour cells, immune effectors, and endothelial cells [5, 6]. Nilsson et al. first demonstrated that the platelet transcriptome is systematically altered by malignancy, showing that platelets harbour tumour-derived RNA biomarkers [7]. This so-called tumour-educated platelet (TEP) reprogramming reflects the RNA uptake, splicing, and processing machinery that platelets inherit from megakaryocyte progenitors, enabling internalisation of tumour-derived RNA cargoes during circulation [8, 7]. Compared to ctDNA, platelets can be isolated in far greater abundance from a standard blood draw and profiled by whole-transcriptome RNA sequencing, yielding signals that reflect both tumour presence and tumour-type-specific molecular biology [9, 10].

Best et al. established the diagnostic potential of TEP RNA sequencing as a multi-class liquid biopsy platform in a landmark Cancer Cell study in 2015 [11]. Profiling whole-blood platelet transcriptomes from 283 blood samples spanning patients with six cancer types - NSCLC, colorectal cancer (CRC), glioblastoma multiforme (GBM), hepatobiliary cancer, breast cancer, and pancreatic cancer alongside healthy donors, they showed that TEP transcriptomes encode cancer-type-specific expression and splicing signatures that support automated tumour-type discrimination [11]. Using an SVM classifier with gene-level feature selection, Best et al. achieved high accuracy for binary cancer-versus-healthy discrimination and meaningful multiclass tumour-type assignment, the first proof-of-concept for blood platelet RNA-seq as a pan-cancer diagnostic tool [11]. The study further demonstrated that the molecular pathways dysregulated in TEPs reflected the established oncogenic biology of the corresponding tumour type, indicating that platelet transcriptomes carry biologically interpretable, cancer-type-specific information rather than non-specific activation noise [11]. The full dataset was deposited in the NCBI GEO under accession GSE68086 [12], enabling independent re-analysis and methodological extension. The original Best et al. study was necessarily constrained in several analytical dimensions. The classification framework evaluated only a single algorithmic family (SVM); the comparative performance of regularised linear classifiers, tree-based ensembles, gradient boosting frameworks, and deep neural networks on GSE68086 had not been examined. Machine learning approaches for transcriptomic cancer classification have advanced substantially since 2015: Random Forests [13], XGBoost [14], and LightGBM [15] consistently perform well in high-dimensional genomic benchmarks [16], and deep learning architectures have expanded the capacity to model complex non-linear feature interactions in large-scale transcriptomic data [17, 18]. Systematic benchmarking has shown repeatedly that no single algorithm dominates across all datasets and class configurations [16], a finding that motivates comprehensive multi-classifier evaluation before any algorithmic recommendation can be translated to a clinical setting. A second gap concerns model interpretability. Predictive accuracy alone is insufficient in clinical diagnostics: predictions must be attributable to specific biological features, enabling biological validation and mechanistic understanding [20]. The opacity of ensemble and deep learning models has been identified as a barrier to their adoption in high-stakes medical decision-making, where clinicians and regulators require explainable predictions [20]. SHAP (SHapley Additive exPlanations), grounded in cooperative game theory [21, 22], decomposes predictions into additive gene-level contributions. Applied to transcriptomic cancer classifiers, class-specific SHAP analysis identifies at single-gene resolution which transcripts most strongly drive classification for each cancer type, generating biologically testable hypotheses and candidate biomarkers for prospective validation [21, 22].

Against this background, four specific gaps remain in the TEP RNA-seq classification literature. First, no comprehensive multi-algorithm benchmarking comparing regularised linear models, kernel methods, tree-based ensembles, gradient boosting, and deep neural networks has been performed on GSE68086. Second, statistical uncertainty in reported classification metrics has not been quantified; percentile bootstrap confidence intervals [23, 24] are essential to determine whether performance differences between classifiers are statistically meaningful given the small test partitions characteristic of this dataset. Third, a complete one-versus-rest differential expression analysis with Benjamini-Hochberg FDR correction [25], coupled with GO and KEGG pathway enrichment analysis [26], has not been systematically applied to characterise cancer-type-specific TEP transcriptomic signatures. Fourth, SHAP-based class-specific feature attribution has not been applied to TEP transcriptomics, leaving single-gene-resolution biological interpretation and candidate biomarker identification unaddressed.

To address these gaps, we undertook a comprehensive transcriptomic and machine learning analysis of GSE68086 [11, 12], specifically: (i) a systematic preprocessing and normalisation pipeline incorporating library-size QC, expression-level filtering, CPM normalisation [27], and MAD-based feature selection to produce a 280 × 5,000 gene feature matrix; (ii) one-versus-rest differential expression analysis (Welch’s t-test with BH-FDR correction [28, 25]) and GO Biological Process 2023 and KEGG 2021 Human pathway enrichment via Enrichr [26]; (iii) systematic benchmarking of seven supervised classifiers - Logistic Regression, SVM (RBF) [29], Random Forest [13], XGBoost [14], LightGBM [15], KNN [30], and an MLP [31, 32, 33] - using stratified 5-fold cross-validation and a held-out test set (scikit-learn [34]); (iv) statistical uncertainty quantification using 2,000-resample percentile bootstrap confidence intervals [23, 24]; and (v) SHAP TreeExplainer [21] applied to the Random Forest classifier for global and cancer-type-specific gene-level attribution rankings. This work makes two complementary contributions. Computationally, it provides the first multi-algorithm benchmarking of TEP RNA-seq classification with bootstrap-based uncertainty quantification, establishing a methodologically transparent and reproducible reference framework for future blood-based transcriptomic cancer classification studies. Translationally, the SHAP-prioritised gene signatures including IFITM3, C6orf62, and ATP5PD - identified consistently across independent classification frameworks represent biologically grounded, experimentally testable candidates for next-generation platelet-based liquid biopsy assays, with direct implications for early cancer detection, tumour-of-origin assignment, and precision oncology.

## 2. Results

### 2.1 Dataset Characteristics and Quality Control

The GSE68086 dataset [12] comprised 280 whole-blood platelet RNA-seq samples representing seven diagnostic categories: lung cancer/NSCLC (n = 60), healthy donors (n = 54), colorectal cancer (CRC; n = 40), glioblastoma multiforme (GBM; n = 40), breast cancer (n = 38), pancreatic cancer (n = 34), and hepatobiliary cancer (n = 14) (Figure 1). Initial parsing of the raw count matrix yielded 57,736 genomic features; after removing entries with missing or ambiguous HGNC symbols and deduplicating - retaining the highest mean-expressing record per symbol - 38,789 unique genes were carried forward. All 280 samples exceeded the minimum library-size threshold of 50,000 reads; none were excluded. Library sizes ranged from 214,812 to 7,124,527 reads (mean ± SD: 2,045,411 ± 1,000,642), with lung cancer and CRC showing greater intra-group variability (Figure 1B). Genes detected per sample (count > 0) ranged from approximately 4,000 to 12,000, with a per-group median of 8,000-9,000 (Figure 1C), consistent with the sparse transcriptomic profiles characteristic of TEP RNA sequencing [11].

**Figure 1.**
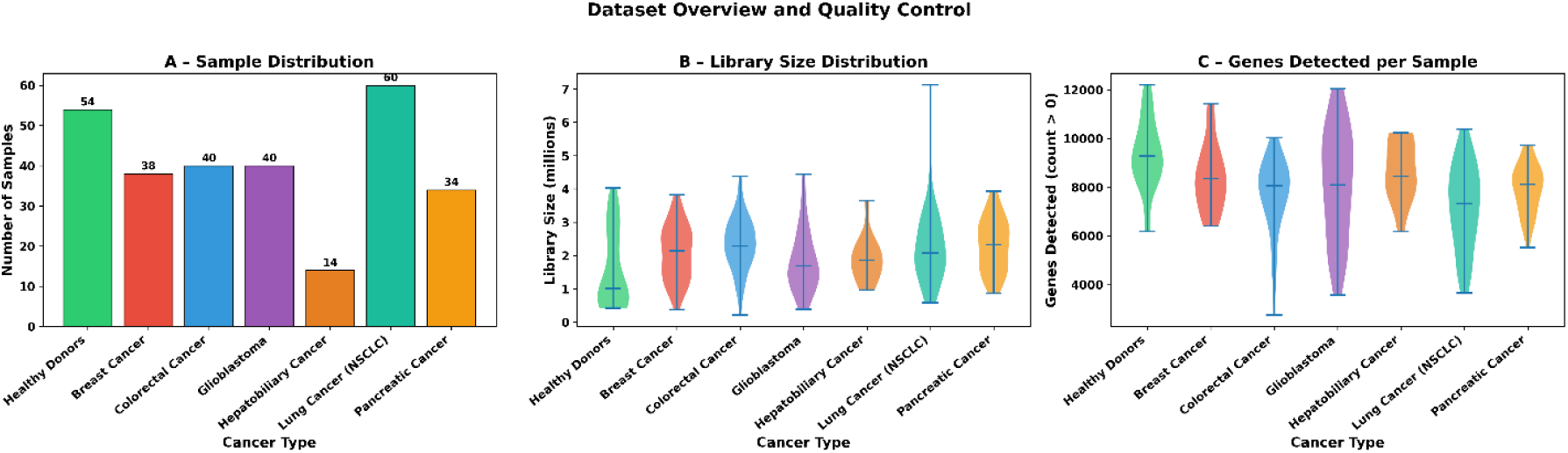
Dataset overview and quality control. (A) Sample counts per diagnostic group. (B) Library-size distributions (millions of reads) stratified by cancer type, displayed as violin plots with median lines. (C) Number of genes detected per sample (count > 0) across groups. NSCLC: non-small cell lung cancer; CRC: colorectal cancer; GBM: glioblastoma multiforme.

### 2.2 Gene Filtering, Normalisation, and Feature Selection

An expression filter retaining genes with ≥ 5 raw counts in at least 10% of samples reduced the feature space from 38,789 to 9,639 genes (75.2% reduction). CPM normalisation followed by log₂(CPM + 1) transformation corrected for inter-sample sequencing depth differences and stabilised count-data variance [27]. Selecting the 5,000 genes with the highest Median Absolute Deviation (MAD) across samples yielded the final 280 × 5,000 feature matrix used in all downstream analyses.

### 2.3 Dimensionality Reduction

PCA, UMAP, and t-SNE were applied to the standardised log₂-CPM feature matrix to characterise global transcriptomic structure (Figure 2). The first two principal components jointly explained 32.5% of total variance (Figure 2A); healthy donor samples showed a partial tendency towards distinct clustering, consistent with their immunologically distinct platelet transcriptome. UMAP revealed greater local structure, with lung cancer samples forming a partially distinct linear arrangement (Figure 2B), while t-SNE provided the clearest visual separation, with healthy donors forming relatively coherent clusters (Figure 2C). The persistent overlap across all three projections indicates that multi-class cancer discrimination from TEP transcriptomes is an inherently difficult problem and motivates the application of supervised machine learning.

**Figure 2.**
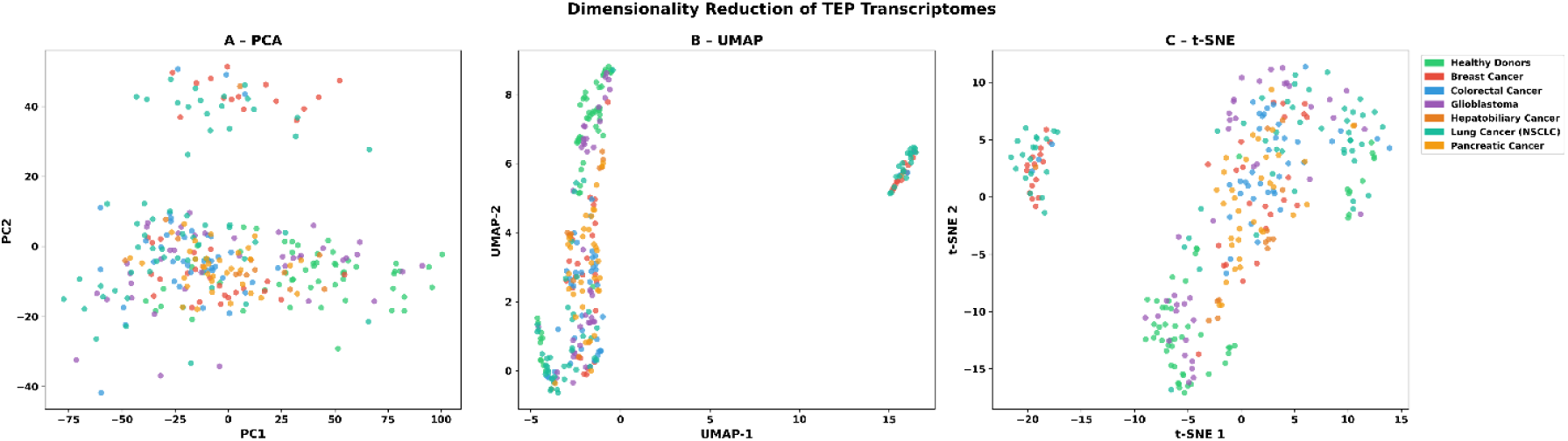
Dimensionality reduction of TEP transcriptomes. (A) Principal Component Analysis (PCA); PC1 and PC2 jointly explain 32.5% of total transcriptomic variance. (B) UMAP projection (n_neighbors = 15, min_dist = 0.1) applied to the 50-component PCA embedding. (C) t-SNE projection (perplexity = 30, 1,000 iterations) applied to the same 50-component PCA embedding. Samples are coloured by diagnostic category; legend is shared across panels.

### 2.4 Differential Expression Analysis

#### 2.4.1 Cancer-Specific Gene Signatures

Welch’s two-sample t-test with BH-FDR correction was applied in a one-versus-rest framework across all 5,000 MAD-selected genes [28, 25]. At thresholds of BH-adjusted q < 0.05 and log_2_FC > 0.5, the number of significantly upregulated genes per class was: healthy donors (3,138), breast cancer (207), GBM (90), CRC (32), and hepatobiliary cancer (17); complete per-class lists are provided in Supplementary Table S1. The substantially larger number of genes upregulated in healthy donors relative to the pooled cancer cohort likely reflects cancer-mediated depletion of immune-cell RNA from TEPs - a hallmark of malignant platelet reprogramming [11].

#### 2.4.2 Expression Heatmap of Top Differentially Expressed Genes

Z-scored log₂-CPM expression of the top ten upregulated genes per class revealed sharply delineated, class-specific expression blocks (Figure 3). Healthy donor samples showed strong upregulation of lymphocyte-associated transcripts, including TTN, IL7R, CD3D, IL2RG, CD3G, and CCR7. Class-specific markers included IGFBP2, NEK7, and DOK6 for breast cancer; HBG2, CRYM, and FECH for CRC; PRUNE2, FKBP5, and MPZL3 for GBM; CD163 and PKP2 for hepatobiliary cancer; FBXW5 and TM4SF1 for NSCLC; and LYZ, NADK, and APOBEC3A for pancreatic cancer.

**Figure 3.**
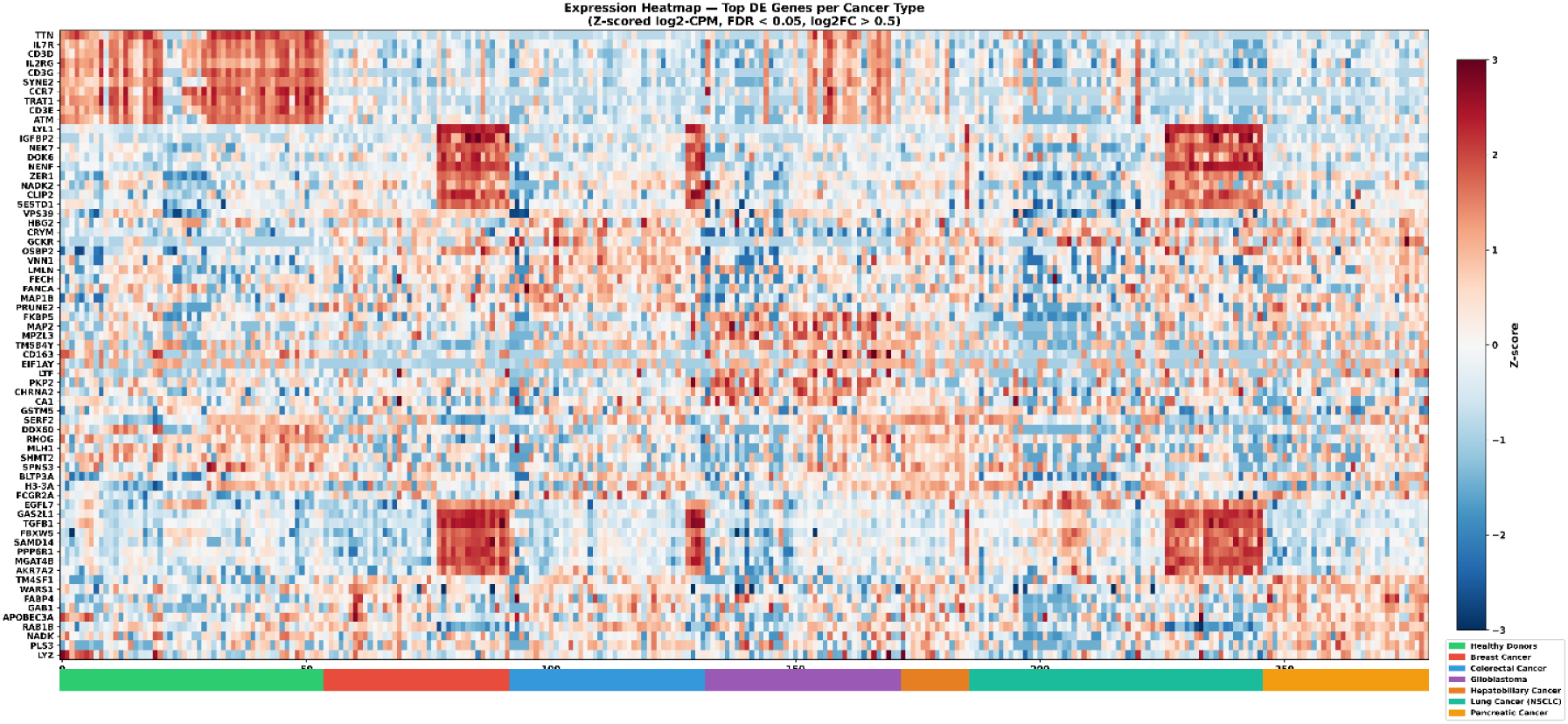
Expression heatmap of the top differentially expressed genes per cancer type. Z-scored log_2_-CPM values for the top ten upregulated genes per class (BH-FDR < 0.05; log_2_FC > 0.5). Samples (columns) are arranged by cancer type as indicated by the colour bar at the bottom. The red-blue diverging colour scale represents Z-scores clipped at ±3.

#### 2.4.3 GO and KEGG Pathway Enrichment

Functional enrichment analysis of significantly upregulated DEGs per cancer type via the Enrichr API [26], querying GO Biological Process 2023 and KEGG 2021 Human gene sets (BH-adjusted p < 0.05), yielded significant results for four cancer types (Figure 4). For CRC, top enriched terms included Oxygen Transport (-log₁₀ p ≈ 4.3), Carbon Dioxide Transport, Gas Transport, Hydrogen Peroxide Catabolic Process, and Positive Regulation of Cell Death. GBM was associated with mucosal and antimicrobial immune response terms (Defence Response to Fungus, Innate Immune Response in Mucosa), likely reflecting defensin-family gene cross-annotation. For NSCLC, the most significant term was Endocytosis (-log₁₀ p ≈ 5.2), followed by Receptor-Mediated Endocytosis, VEGF Signalling Pathway, and the directly corroborating KEGG term Non-Small Cell Lung Cancer. Pancreatic cancer yielded enrichment of Actin Filament Network Formation and Regulation of Protein Transport. Breast cancer and hepatobiliary cancer did not yield significant enrichment terms, attributable to small DEG sets (207 and 17 genes, respectively) and molecular heterogeneity within these groups.

**Figure 4.**
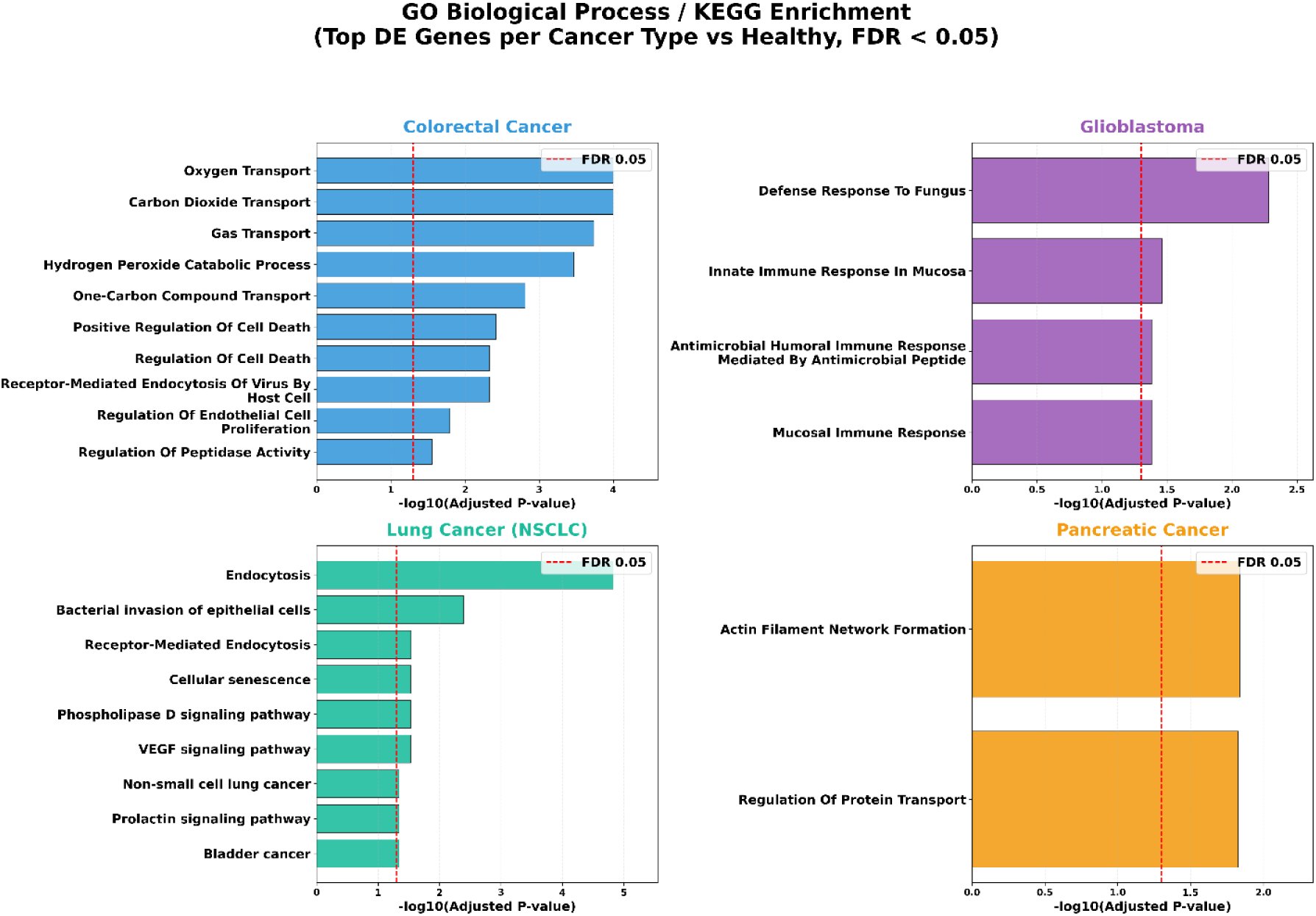
GO Biological Process and KEGG pathway enrichment for four cancer types. Bars represent -log_10_(adjusted p-value); the red dashed vertical line indicates the FDR = 0.05 significance threshold. Enrichment was performed using Enrichr [26] on all statistically significant upregulated DEGs per cancer type. Only cancer types with at least one term surviving BH adjustment are shown.

### 2.5 Machine Learning Classification

#### 2.5.1 Comparative Model Performance

Six supervised machine learning classifiers and one deep learning model were trained on the 5,000-gene log₂-CPM feature matrix using an 80/20 stratified train-test split (224 training / 56 test samples) with stratified 5-fold cross-validation on the training set. Performance metrics are summarised in Table 1.

**Table 1.** Classification performance of all models on the held-out test set (n = 56). Models are ranked by test F1-macro (descending). CV: 5-fold stratified cross-validation on training data; F1: macro-averaged F1-score; Prec: macro-averaged precision; Rec: macro-averaged recall; AUC: one-vs-rest macro-averaged ROC-AUC; SD: standard deviation; ML: classical machine learning; DL: deep learning. Bold values denote best performance per metric.

| Model | Type | CV Acc<br>(Mean±S<br>D) | CV F1<br>(Mean±S<br>D) | Test Acc | Test F1 | Prec | Rec | AUC |
| --- | --- | --- | --- | --- | --- | --- | --- | --- |
| Logistic Regression | ML | 0.589±0.079 | 0.515±0.082 | 0.518 | 0.522 | 0.532 | 0.546 | 0.869 |
| SVM (RBF) | ML | 0.554±0.070 | 0.481±0.043 | 0.482 | 0.471 | 0.531 | 0.460 | 0.865 |
| XGBoost | ML | 0.580±0.086 | 0.514±0.087 | 0.482 | 0.469 | 0.567 | 0.448 | 0.828 |
| Random Forest | ML | 0.518±0.069 | 0.424±0.061 | 0.464 | 0.464 | 0.561 | 0.441 | 0.823 |
| MLP | DL | - | - | 0.446 | 0.435 | 0.441 | 0.466 | 0.866 |
| LightGBM | ML | 0.549±0.088 | 0.464±0.057 | 0.429 | 0.364 | 0.392 | 0.364 | 0.816 |
| KNN | ML | 0.393±0.066 | 0.321±0.058 | 0.357 | 0.346 | 0.477 | 0.347 | 0.723 |

Logistic Regression achieved the highest test macro F1-score (0.522) and ROC-AUC (0.869), along with the best cross-validated accuracy (0.589 ± 0.079) and F1-macro (0.515 ± 0.082), indicating consistent generalisation across training folds. SVM (RBF) ranked second by test F1 (0.471) with a closely comparable AUC (0.865). XGBoost achieved the highest test precision (0.567) despite a test F1 of 0.469. LightGBM and KNN were the weakest classical classifiers (F1: 0.364 and 0.346, respectively), with KNN recording the lowest AUC (0.723) - reflecting its sensitivity to high-dimensional feature spaces [30]. MLP cross-validation was not performed; training used early stopping on an internal 15% validation split.

#### 2.5.2 Best Model: Confusion Matrix and ROC Curve Analysis

Detailed performance of the best-performing model (Logistic Regression) is shown in Figure 5. The normalised confusion matrix (Figure 5A) revealed marked inter-class differences in recall. Hepatobiliary cancer achieved perfect recall (1.00; 3/3 test samples); healthy donors reached 0.82 (9/11 correctly classified); pancreatic cancer and GBM reached 0.57 and 0.50, respectively. Breast cancer showed the lowest recall (0.14; 1/7 correctly classified), with 71% of samples misclassified as pancreatic cancer - suggesting substantial transcriptomic overlap between these two classes in the TEP feature space. Confusion matrices for all six classical models are provided in Supplementary Figure S1. Per-class ROC curves (Figure 5B) yielded AUC values of 1.000 (hepatobiliary), 0.978 (healthy), 0.904 (GBM), 0.851 (breast), 0.848 (pancreatic), 0.790 (lung), and 0.714 (CRC), giving a macro-average of 0.869 [35]. The high AUC values relative to moderate hard-classification accuracy indicate well-calibrated posterior probabilities even when the argmax class assignment is incorrect.

**Figure 5.**
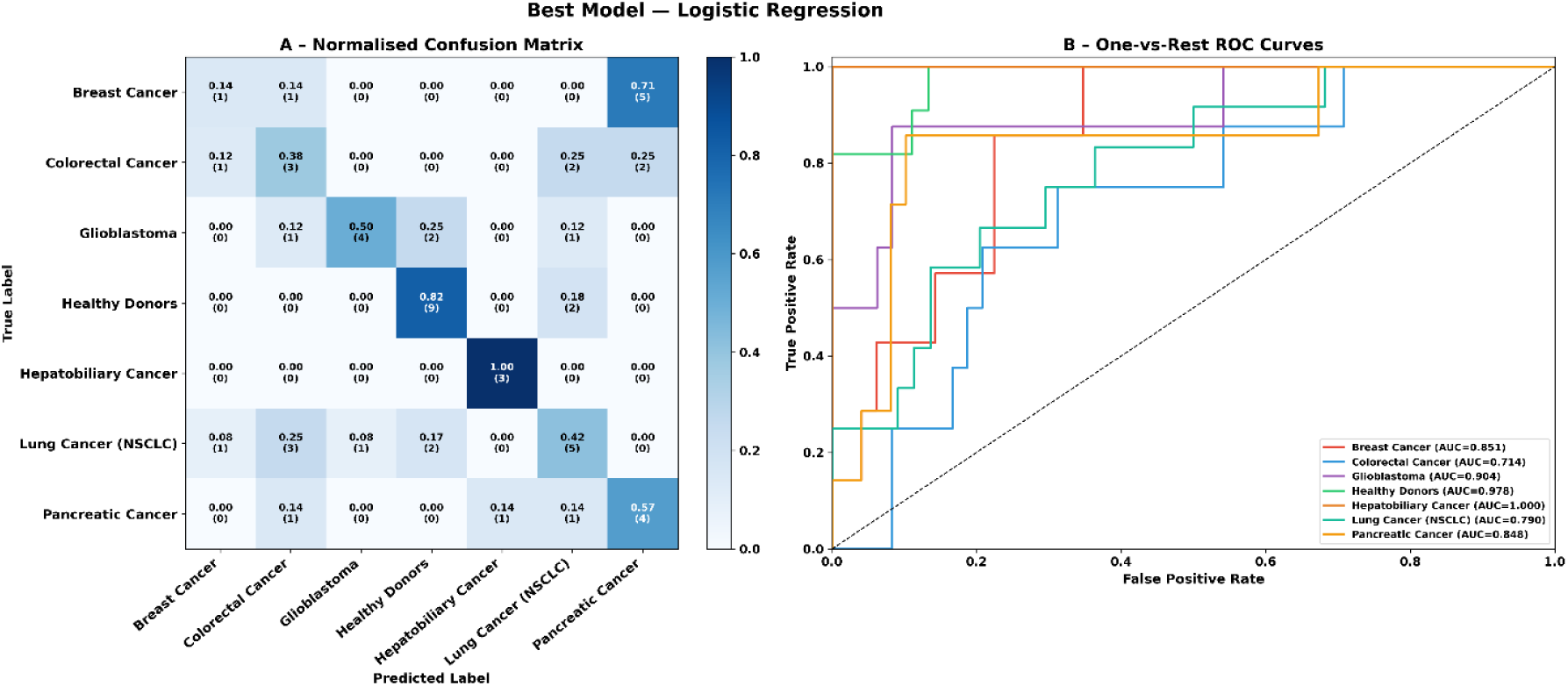
Performance of the best-performing model (Logistic Regression; test F1 = 0.522; macro-AUC = 0.869). (A) Normalised confusion matrix; cell values represent the fraction of true-class samples predicted to each class; raw counts are shown in parentheses. (B) One-vs-rest ROC curves per cancer type with individual AUC values.

### 2.6 Deep Learning Classification: Multilayer Perceptron

The MLP achieved a test accuracy of 0.446, macro F1 of 0.435, and AUC of 0.866 (Table 1). Although its AUC matched the top classical models, hard-label metrics fell below Logistic Regression, SVM, and XGBoost. Training curves (Supplementary Figure S2) showed marked overfitting: training accuracy approached ∼100% by approximately epoch 45 while validation accuracy plateaued at 65-70%, with a widening loss gap despite Dropout regularisation (rates: 0.50/0.50/0.30) and L2 weight decay (λ = 10⁻⁴) [33]. Early stopping with patience = 20 epochs terminated training at epoch 45, restoring minimum-validation-loss weights. The MLP confusion matrix and per-class ROC curves are provided in Supplementary Figure S3.

### 2.7 Bootstrap Confidence Intervals

To quantify statistical uncertainty on the held-out test set (n = 56), 95% percentile bootstrap confidence intervals were computed for all five performance metrics using 2,000 resamples drawn with replacement (seed = 42) [23, 24]. Results are summarised in Table 2 and visualised as a forest plot in Figure 6.

**Figure 6.**
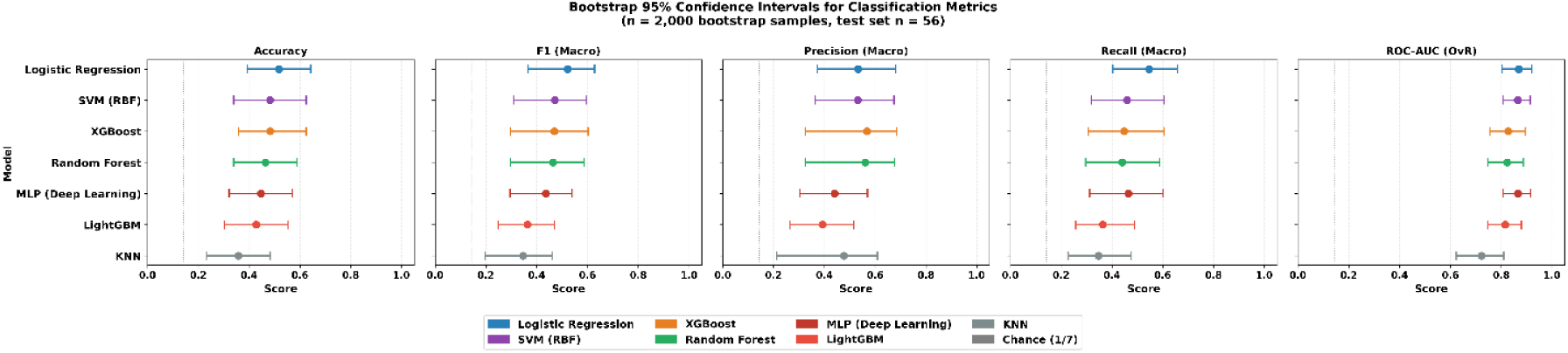
Bootstrap 95% confidence interval forest plot for five classification metrics. Each filled circle represents the point estimate; horizontal bars represent the 95% percentile CI (n = 2,000; test set n = 56). Models are ordered by F1-macro point estimate, with the best-performing model at the top. The vertical grey dashed line indicates the chance level (1/7 ≈ 0.14).

**Table 2.** Bootstrap 95% confidence intervals for all classifiers. Point estimate [95% CI lower-upper]. n = 2,000 resamples; test set n = 56. All metrics are macro-averaged across seven classes. Models are ranked by F1-macro point estimate. Bold denotes highest point estimate per metric.

| Model | Accuracy | F1 (Macro) | Precision | Recall | ROC-AUC |
| --- | --- | --- | --- | --- | --- |
| Logistic Regression | 0.518<br>[0.393-0.643] | 0.522<br>[0.367-0.628] | 0.533<br>[0.372-0.682] | 0.546<br>[0.404-0.659] | 0.869<br>[0.804-0.924] |
| SVM (RBF) | 0.482<br>[0.339-0.625] | 0.471<br>[0.309-0.596] | 0.531<br>[0.364-0.674] | 0.460<br>[0.319-0.605] | 0.865<br>[0.802-0.924] |
| XGBoost | 0.482<br>[0.357-0.625] | 0.469<br>[0.297-0.602] | 0.567<br>[0.326-0.685] | 0.448<br>[0.308-0.606] | 0.828<br>[0.754-0.898] |
| Random Forest | 0.464<br>[0.339-0.589] | 0.464<br>[0.297-0.585] | 0.561<br>[0.326-0.678] | 0.441<br>[0.296-0.588] | 0.823<br>[0.744-0.898] |
| MLP | 0.446<br>[0.321-0.571] | 0.435<br>[0.294-0.540] | 0.441<br>[0.304-0.570] | 0.466<br>[0.313-0.602] | 0.866<br>[0.805-0.924] |
| LightGBM | 0.429<br>[0.304-0.554] | 0.364<br>[0.248-0.470] | 0.392<br>[0.265-0.517] | 0.364<br>[0.257-0.488] | 0.816<br>[0.741-0.888] |
| KNN | 0.357<br>[0.232-0.482] | 0.346<br>[0.196-0.461] | 0.477<br>[0.214-0.609] | 0.347<br>[0.227-0.476] | 0.723<br>[0.621-0.810] |

The wide confidence intervals for accuracy and F1 (typical CI width ≈ 0.25) reflect the small test-set size (n = 56) rather than model instability. Two findings, however, hold across all models. AUC CIs are substantially narrower than hard-label metric CIs (typical width ≈ 0.12), confirming that probability-based discriminability is estimated more reliably than hard-label accuracy in small-sample settings. The only pair-wise comparison with statistical support is Logistic Regression versus KNN: Logistic Regression’s AUC lower bound (0.804) does not overlap KNN’s upper bound (0.810). For all other metric-model combinations, overlapping CIs indicate that observed ranking differences may reflect sampling variability rather than true model superiority, underscoring the need for larger independent validation cohorts.

### 2.8 SHAP Feature Importance Analysis

#### 2.8.1 Global SHAP Importance

SHAP values were computed for the Random Forest classifier using TreeExplainer on the 56 test samples, yielding a 56 × 5,000 × 7 attribution tensor [21]. Global feature importance - the mean absolute SHAP value averaged across all test samples and seven cancer classes - identified IFITM3 as the most influential gene (|ϕ| = 3.17 × 10^-3^), followed by C6orf62 (2.17 × 10^-3^), VPS13C (1.86 × 10^-3^), LINC00861 (1.82 × 10^-3^), LUC7L (1.77 × 10^-3^), MDM4 (1.70 × 10^-3^), and ATP5PD (1.66 × 10^-3^) (Figure 7).

**Figure 7.**
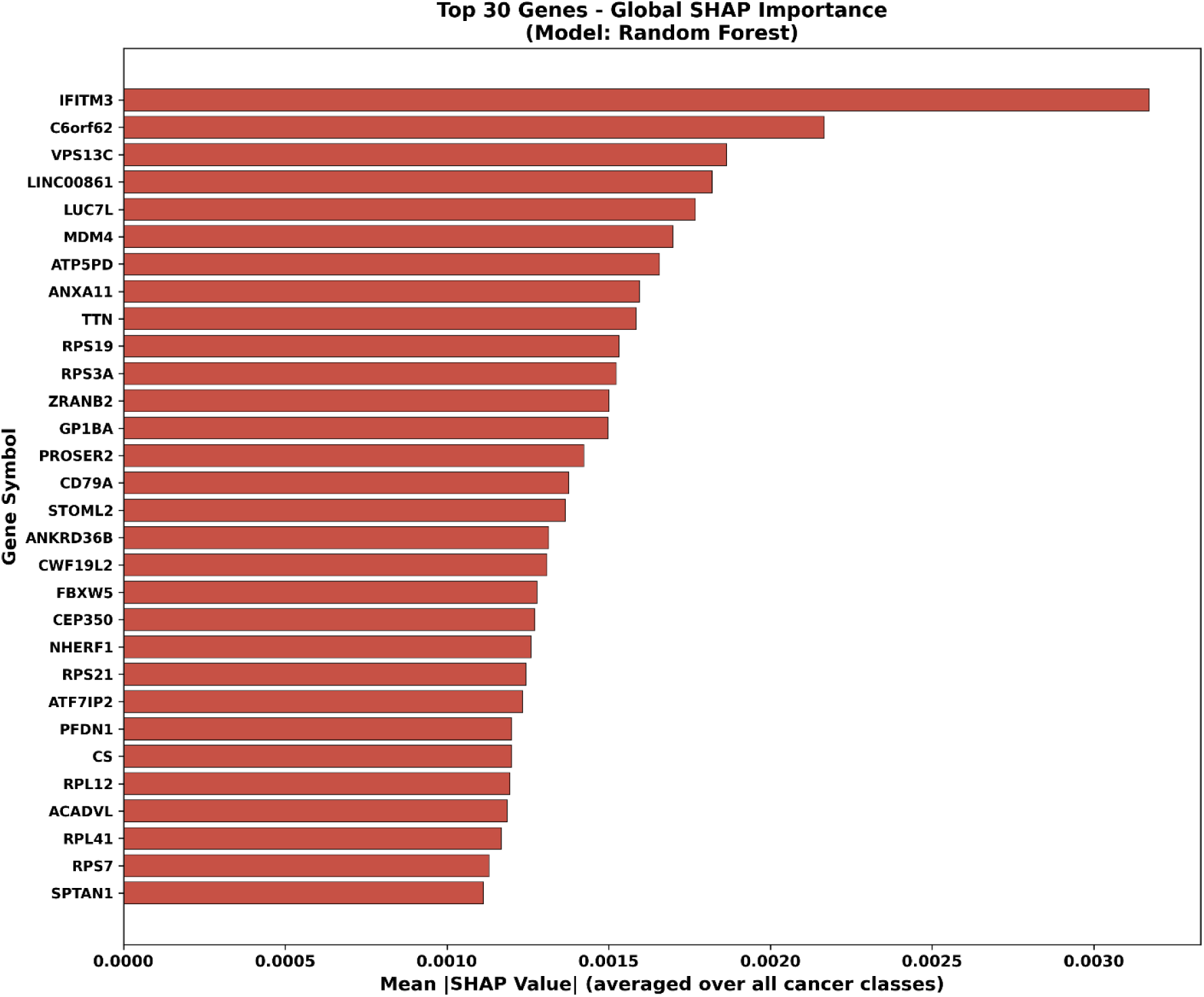
Global SHAP feature importance - top 30 genes (Random Forest classifier). Mean absolute SHAP value (|ϕ|) for each gene, averaged over all 56 test samples and all seven cancer classes. Higher values indicate greater average contribution to classification decisions irrespective of cancer type.

#### 2.8.2 Cancer-Type-Specific SHAP Analysis

Per-class SHAP analysis revealed distinct discriminative gene sets for each cancer type (Figure 8).

- **Healthy donors:** VPS13C (|ϕ| = 5.07 × 10^-3^), RPS3A (4.84 × 10^-3^), and LINC00861 (4.74 × 10^-3^) were the leading discriminators, consistent with the immune-rich transcriptome of non-malignant platelets.
- **Breast cancer:** TMSB4Y (2.17 × 10^-3^), EVI2A (1.86 × 10^-3^), and PROSER2 (1.74 × 10^-3^) were the top contributors.
- **CRC:** IFITM3 (3.73 × 10^-3^), ACADVL (1.99 × 10^-3^), and C6orf62 (1.77 × 10^-3^) were most discriminative.
- **GBM:** IFITM3 was overwhelmingly dominant (7.37 × 10^-3^), with NHERF1 (2.90 × 10^-3^) and CEP350 (2.73 × 10^-3^) as secondary contributors.
- **Hepatobiliary cancer:** ATP5PD (4.64 × 10^-3^), STOML2 (3.78 × 10^-3^), and PFDN1 (3.27 × 10^-3^) indicated a mitochondrial and proteostasis-related signature.
- **NSCLC:** C6orf62 (4.23 × 10^-3^), ANXA11 (4.09 × 10^-3^), and CS (citrate synthase; 3.83 × 10^-3^) were the top features, the latter reflecting TCA cycle activity in NSCLC-TEPs.
- **Pancreatic cancer:** C6orf62 (4.35 × 10^-3^) and IFITM3 (3.49 × 10^-3^) dominated, while PROSER2 (2.31 × 10^-3^) and GRAMD1A (2.22 × 10^-3^) provided type-specific signals.

**Figure 8.**
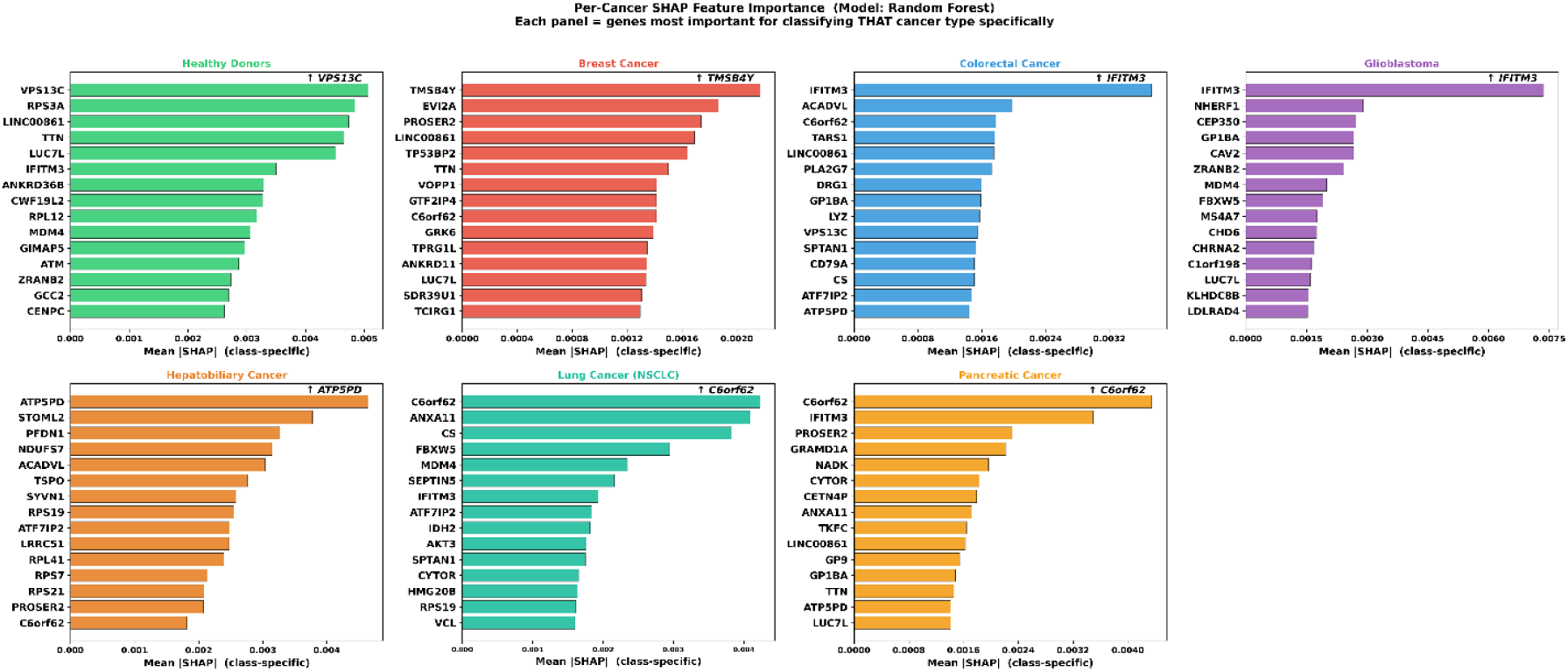
Per-cancer-type SHAP feature importance - top 15 genes per class (Random Forest classifier). Each panel shows the top 15 genes ranked by mean class-specific absolute SHAP value. Panel colour corresponds to the cancer-type colour scheme used throughout. Genes appearing across multiple panels represent shared discriminative features; cancer-exclusive genes reflect type-specific platelet transcriptomic reprogramming signals.

**Figure 9:**
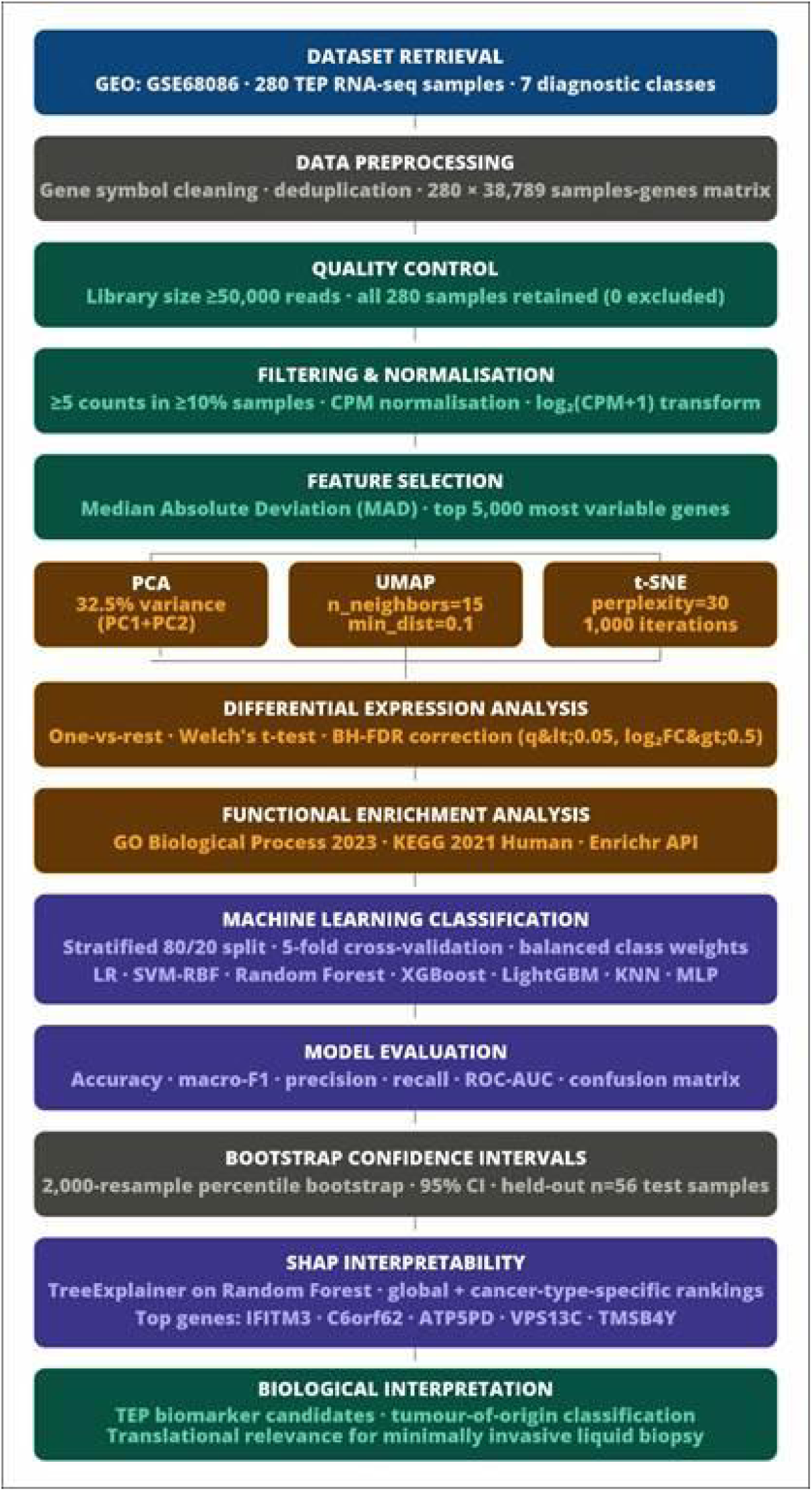
Overview of the transcriptomic preprocessing, differential expression, machine learning classification, and SHAP-based interpretability workflow for pan-cancer prediction using tumor-educated platelet (TEP) RNA-seq data from GSE68086.

IFITM3 ranked first in Random Forest (importance = 4.58 × 10^-3^); MPL led in XGBoost (9.95 × 10^-3^); FKBP5 led in LightGBM (score = 55). These rankings corroborate the SHAP global analysis in identifying IFITM3 as a consistently important pan-cancer TEP discriminator.

## 3. Methods

### 3.1 Dataset Retrieval and Study Design

The GSE68086 dataset [12] was obtained from the NCBI Gene Expression Omnibus (GEO). Originally described by Best et al. [11], it comprises RNA-seq profiles from tumour-educated blood platelets (TEPs) collected from patients with seven clinical conditions: breast cancer, CRC, GBM, hepatobiliary carcinoma, NSCLC, pancreatic adenocarcinoma, and a healthy donor cohort. TEPs are a biologically accessible transcriptomic source: platelets sequester and preserve tumour-derived RNA, enabling minimally invasive molecular profiling of systemic malignancies [11]. This study extends the original classification framework of Best et al. [11] which used SVM with leave-one-out cross-validation by benchmarking a broader suite of machine learning and deep learning approaches alongside differential expression and pathway analyses.

The complete raw gene expression count matrix was downloaded directly from GEO. The matrix comprised 57,738 rows (including two header rows) and 282 columns (including two annotation columns), encoding read counts for 57,736 genomic features across 280 patient samples. Sample-level metadata, including cancer-type annotations and sample identifiers, were embedded in the first two rows and parsed accordingly. The dataset comprised the following class distribution: healthy donors (n = 54), lung cancer/NSCLC (n = 60), CRC (n = 40), GBM (n = 40), breast cancer (n = 38), pancreatic cancer (n = 34), and hepatobiliary cancer (n = 14). All analyses were implemented in Python (version 3.10.0) and executed within the Google Colaboratory cloud computing environment.

### 3.2 Data Preprocessing and Cleaning

The raw count matrix was loaded using the pandas library with tab-delimited parsing and no assumed header row. The structural layout placed HGNC gene symbols in the second column and raw integer read counts in all subsequent columns. Count values were coerced to numeric format, with non-numeric entries converted to zero via pd.to_numeric(errors=‘coerce’), followed by zero-filling of any residual missing values. The resulting expression array was cast to 32-bit floating-point representation to optimize memory efficiency. Gene symbol annotation was extracted from the second column. Entries corresponding to missing, null, or placeholder gene identifiers were removed before further processing. The source dataset contained multiple genomic features annotated with identical HGNC gene symbols, arising from alternative transcripts, pseudogenes, or annotation redundancy; a deduplication strategy was therefore applied. For each duplicated gene symbol, the mean expression value across all samples was computed, and only the feature with the highest mean expression was retained. This ensures the most transcriptionally active isoform is preferentially selected. Following deduplication, 38,789 unique gene symbols remained, and the matrix was transposed to the conventional samples-by-features orientation (280 samples × 38,789 genes).

### 3.3 Quality Control

Sample-level QC was performed by computing total library size (sum of all counts) and the number of detected genes (count > 0) for each sample. A minimum library-size threshold of 50,000 total reads was applied to exclude samples with severely under-sequenced profiles. No samples in GSE68086 fell below this threshold (minimum observed: 214,812 reads; mean ± SD: 2,045,411 ± 1,000,642; maximum: 7,124,527); all 280 samples were retained. Library size distributions and gene detection counts were visualised as violin plots stratified by cancer type, to assess sequencing depth homogeneity across diagnostic groups.

### 3.4 Gene Filtering, Normalisation, and Feature Selection

#### 3.4.1 Expression-Based Gene Filtering

To remove lowly expressed and non-informative genes, a minimum expression filter was applied: a gene was retained only if it exhibited a raw count of ≥ 5 in at least 10% of the total sample cohort (a minimum of 28 samples). This threshold mirrors filtering heuristics common in bulk RNA-seq pipelines and mitigates the influence of dropout events and background noise on downstream statistical inference [27]. Following this filter, 9,639 genes were retained from the original 38,789, representing a ∼75% reduction in feature dimensionality.

#### 3.4.2 Library Size Normalisation and Log-Transformation

Raw count data were normalised to counts per million (CPM) by dividing each sample’s per-gene count by the sample’s total library size and multiplying by 10⁶. This step corrects for inter-sample differences in sequencing depth while preserving proportional gene expression levels. Normalised CPM values were subsequently subjected to a log₂(CPM + 1) transformation to stabilise the variance-mean relationship inherent to count data and reduce the influence of highly expressed outlier genes. A pseudocount of 1 was added before log-transformation to prevent undefined values at zero counts [27].

#### 3.4.3 High-Variance Gene Selection via Median Absolute Deviation

Following normalisation, a gene-level variability filter was applied to reduce dimensionality while retaining the most biologically informative features. For each gene, the Median Absolute Deviation (MAD) was computed across all 280 samples in the log₂(CPM + 1) space. MAD is a robust dispersion measure less sensitive to outliers than standard deviation, making it well-suited to high-dimensional transcriptomic data. The 5,000 genes with the highest MAD values were selected, yielding a final feature matrix of 280 × 5,000 used in all subsequent analyses.

### 3.5 Exploratory Data Analysis and Dimensionality Reduction

PCA [36, 37], UMAP [38], and t-SNE [39] were applied to the standardised log_2_-CPM feature matrix to characterise global transcriptomic structure and assess inter-class separability. The 5,000-gene matrix was first scaled to zero mean and unit variance using StandardScaler (scikit-learn [34]). PCA was performed retaining the first 50 principal components (random_state = 42); the first two jointly explained 32.5% of total transcriptomic variance. UMAP was applied to the 50-component PCA embedding with parameters n_components = 2, n_neighbors = 15, min_dist = 0.1, and random_state = 42. t-SNE was independently applied to the same 50-component PCA embedding using a perplexity of 30, 1,000 optimisation iterations, and random_state = 42. Two-dimensional scatter plots were generated with samples colour-coded by cancer type.

### 3.6 Differential Expression Analysis

#### 3.6.1 Cancer-Specific Gene Signatures

To identify cancer-type-specific gene expression signatures, a one-versus-rest differential expression framework was implemented. For each of the seven diagnostic groups, Welch’s two-sample t-test (ttest_ind with equal_var = False; scipy.stats) was applied independently to all 5,000 genes in the log₂(CPM + 1) feature space [28]. This test does not assume homogeneity of variance between groups. log₂ fold-change (log₂FC) was computed as the difference between mean log_2_(CPM + 1) values of the group of interest and the complement group.

Raw p-values were adjusted using the Benjamini-Hochberg (BH) procedure [25] (statsmodels.stats.multitest.multipletests) to control the false discovery rate across 5,000 simultaneous hypothesis tests. Genes were considered significantly upregulated in the group of interest if they satisfied both: (i) BH-adjusted q < 0.05, and (ii) log₂FC > 0.5. Cancer-specific upregulated gene signatures were ranked by descending fold-change, and the top 20 and top 50 genes per group were exported for downstream functional annotation.

#### 3.6.2 Pan-Cancer Gene Identification

To identify genes consistently upregulated across all malignant classes relative to healthy donors, a pan-cancer analysis was performed. For each of the six cancer types, genes significantly upregulated versus the healthy donor group were determined using the same Welch t-test and BH-FDR framework (q < 0.05; log₂FC > 0.5). The intersection of these six cancer-specific significant gene sets was then computed; genes in this intersection represent transcriptomic features consistently elevated across all cancer types relative to the non-malignant baseline.

#### 3.6.3 Heatmap Visualisation

An expression heatmap was generated to visualise the differential expression landscape of the top 10 upregulated genes per cancer type. Expression values in the log_2_(CPM + 1) space were Z-scored across samples to facilitate cross-gene comparisons. Samples were arranged in cancer-type-sorted order, and the matrix was rendered using a diverging red-blue colour map (RdBu_r) with Z-score limits of ±3.

### 3.7 Machine Learning Classification Pipeline

#### 3.7.1 Data Partitioning and Class Imbalance Handling

The log_2_(CPM + 1) normalised, top-5,000-MAD-gene feature matrix served as input to all classifiers. Class labels were integer-encoded using LabelEncoder. The dataset was partitioned into training (80%) and held-out test (20%) subsets using stratified random sampling (train_test_split, stratify = y, random_state = 42), yielding 224 training and 56 test samples. Stratification preserved the proportion of each cancer type in both partitions. To address class imbalance during training, balanced class weights were computed using scikit-learn’s compute_class_weight(‘balanced’) utility.

#### 3.7.2 Feature Preprocessing within Classifier Pipelines

All classical machine learning models were embedded in scikit-learn Pipeline objects to ensure consistent preprocessing within cross-validation folds and prevent data leakage. Two pipeline architectures were employed: (i) for tree-based ensemble methods (Random Forest, XGBoost, LightGBM), the pipeline consisted of feature standardisation via StandardScaler followed directly by the classifier; (ii) for linear and distance-based methods (SVM, Logistic Regression, k-NN), an additional univariate feature selection step was incorporated using SelectKBest with the ANOVA F-statistic (f_classif) computed on training data only. The number of features retained by SelectKBest was set to 1,500 for SVM and Logistic Regression, and to 500 for k-NN.

#### 3.7.3 Classifier Specifications and Hyperparameters

Six classical machine learning classifiers were evaluated with the following configurations:

- Random Forest [13]: n_estimators = 100, max_features = ‘sqrt’, class_weight = ‘balanced’, random_state = 42, n_jobs = -1.
- SVM-RBF [29]: C = 10, gamma = ‘scale’, class_weight = ‘balanced’, probability estimates enabled (Platt scaling), random_state = 42.
- Logistic Regression (Multinomial): C = 1.0, solver = ‘saga’, maximum iterations = 2,000, multi_class = ‘multinomial’, class_weight = ‘balanced’, random_state = 42.
- XGBoost [14]: n_estimators = 100, max_depth = 4, learning_rate = 0.1, subsample = 0.8, colsample_bytree = 0.8, eval_metric = ‘mlogloss’, random_state = 42.
- LightGBM [15]: n_estimators = 100, num_leaves = 31, learning_rate = 0.1, class_weight = ‘balanced’, random_state = 42.
- k-NN [30]: n_neighbors = 7, distance metric = Euclidean.

#### 3.7.4 Cross-Validation and Test-Set Evaluation

All six classifiers were evaluated using stratified 5-fold cross-validation (StratifiedKFold, n_splits = 5, shuffle = True, random_state = 42) applied exclusively to the training set. Cross-validated accuracy and macro-averaged F1-score (mean ± SD across folds) were reported. Each model was subsequently retrained on the complete training partition and evaluated on the held-out test set. The following metrics were computed on the test set: overall accuracy, macro-averaged precision, macro-averaged recall, macro-averaged F1-score, and multi-class ROC-AUC via the one-versus-rest strategy with macro-averaging [35]. Per-class ROC curves and normalised confusion matrices were generated for all classifiers.

#### 3.7.5 Bootstrap Estimation of Confidence Intervals

Bootstrap confidence intervals were computed for each classifier using the percentile bootstrap method [23, 24] to quantify statistical uncertainty in all reported performance metrics. A total of 2,000 bootstrap resamples were drawn with replacement from the 56 test samples. For each resample, all five performance metrics were recomputed for every model using the corresponding stored predictions and predicted probabilities. Bootstrap samples in which fewer than two distinct classes were represented were discarded from the AUC computation to avoid undefined values. The 2.5th and 97.5th percentiles of the resulting empirical distributions were taken as the lower and upper bounds of the 95% confidence interval for each metric-model combination. A fixed random seed of 42 was used throughout to ensure reproducibility [24], with resampling implemented using numpy.random.default_rng.

### 3.8 Deep Learning Classification: Multilayer Perceptron

A feed-forward multilayer perceptron (MLP) was constructed using TensorFlow 2.20.0 [31] and the Keras functional API. The input layer accepted the 5,000-dimensional log₂(CPM + 1) feature vector following zero-mean unit-variance standardisation (fitted on training data only). The network comprised three fully connected hidden layers of decreasing width: 512 → 256 → 128 neurons. Each hidden layer was followed sequentially by Batch Normalisation [32], a ReLU activation, and a Dropout layer [33] with rates of 0.50, 0.50, and 0.30, respectively. ℓ₂ weight regularisation (λ = 1 × 10⁻⁴) was applied to the kernel weights of each hidden dense layer. The output layer comprised seven neurons with softmax activation.

The model was compiled with the Adam optimiser [40] at an initial learning rate of 1 × 10^-3^, using categorical cross-entropy as the loss function. Training ran for a maximum of 200 epochs with a mini-batch size of 32; 15% of the training data was withheld for monitoring. Sample weights derived from class frequency inversions were applied to address class imbalance. Two callbacks were employed: (i) EarlyStopping with patience = 20 monitoring validation loss, restoring best-weight parameters upon termination; and (ii) ReduceLROnPlateau, which reduced the learning rate by a factor of 0.5 after 10 consecutive epochs of non-improvement, with a minimum allowable rate of 1 × 10^-6^.

### 3.9 Feature Importance and Explainability Analysis

#### 3.9.1 Tree-Based Feature Importances

For tree-based ensemble classifiers (Random Forest, XGBoost, LightGBM), built-in feature importance scores were extracted via the feature_importances_ attribute, reflecting mean decrease in impurity (Gini-based, for Random Forest) or gradient-based gain (for XGBoost and LightGBM) attributable to each feature across all decision trees. The top 20 genes by feature importance for each model were exported as tabular summaries.

#### 3.9.2 SHAP Analysis

SHAP values [21] were computed using shap.TreeExplainer, which leverages the tree structure to compute exact Shapley values in polynomial time. SHAP analysis was applied to the Random Forest classifier as the primary interpretability target. TreeExplainer was fitted on the scaled test feature matrix, producing a SHAP value tensor of shape (56 × 5,000 × 7) (test samples × genes × cancer classes).

Global feature importance was derived by computing the mean absolute SHAP value for each gene, averaged across all test samples and seven cancer classes, capturing each gene’s average marginal contribution to classification decisions. Class-specific feature importance was derived by extracting the SHAP slice for each cancer type and computing the mean absolute SHAP value per gene across test samples. A cross-cancer SHAP heatmap was constructed from the union of the top 10 genes per class (up to 25 genes total), displaying mean absolute class-specific SHAP values in a genes-by-cancer-type matrix.

### 3.10 Gene Ontology and Pathway Enrichment Analysis

Functional annotation of cancer-specific transcriptomic signatures was performed using the Enrichr API [26], accessed via the gseapy Python library [10]. For each cancer type (excluding the healthy donor group), all statistically significant upregulated genes (BH-FDR < 0.05; log_2_FC > 0.5) were submitted to Enrichr. Two gene set libraries were queried: GO Biological Process 2023 and KEGG 2021 Human.

Enrichment results were filtered to exclude terms associated with infectious, parasitic, or viral pathology (e.g., malaria, tuberculosis, HIV/AIDS), which can arise as statistical artefacts from shared immune gene signatures in platelet transcriptomes. Terms were considered significant if the BH-adjusted p-value was < 0.05. The top ten terms per cancer type were visualised as horizontal bar plots with -log_10_(adjusted p-value) along the horizontal axis.

### 3.11 Statistical Analysis

Differential expression analysis was based on Welch’s two-sample t-test, selected for its robustness under unequal sample sizes and heterogeneous variances [28]. Multiple testing correction used the Benjamini-Hochberg method [25] at a 5% FDR threshold. Statistical analyses were implemented using scipy.stats and statsmodels. Machine learning performance metrics were computed using sklearn.metrics [34]. Multi-class ROC-AUC was computed via the one-versus-rest strategy with macro-averaging [35]. Bootstrap confidence intervals were estimated using 2,000 percentile-bootstrap resamples of the held-out test set (Section 3.7.5) [23, 24]. A fixed random state (SEED = 42) was applied throughout to ensure full reproducibility.

### 3.12 Software and Computational Environment

All analyses were implemented in Python (version 3.10.0) and executed in the Google Colaboratory cloud environment. The following libraries were used: pandas and numpy for data handling and numerical computation; scipy.stats and statsmodels for statistical analysis; scikit-learn [34] for machine learning pipelines, classifiers, preprocessing, feature selection, and metrics; XGBoost [14] and lightgbm [15] for gradient boosting; tensorflow 2.20.0/keras [31] for deep learning; shap [21] for explainability; umap-learn [38] and scikit-learn TSNE [39] for dimensionality reduction; matplotlib and seaborn for visualisation; and gseapy [10] for functional enrichment analysis. All figures were generated at 300 DPI resolution.

## 4. Discussion

### 4.1 TEP Transcriptomics as a Multi-Class Liquid Biopsy Platform

The central finding of this study is that TEP RNA-seq profiles from the GSE68086 dataset can support automated seven-class cancer discrimination, with Logistic Regression achieving a macro F1-score of 0.522 and AUC of 0.869 on the held-out test set. These results extend the seminal work of Best et al. [11], who first demonstrated TEP-based diagnostic utility using binary and pan-cancer SVM classification by simultaneously benchmarking seven diverse classifier architectures and integrating differential expression analysis, SHAP-based explainability, and GO/KEGG pathway annotation into a single reproducible pipeline. All 280 samples passed QC, with consistent library size and gene detection profiles across all seven groups, confirming the technical quality of the GSE68086 transcriptomes as a reliable benchmark for pan-cancer classification studies.

### 4.2 Feature Space and Dimensionality Reduction

Reducing from 57,736 raw features to 5,000 MAD-selected genes is a necessary step to manage the curse of dimensionality inherent in small-sample transcriptomic machine learning. CPM normalisation with log₂ transformation is standard for correcting inter-sample depth differences in bulk RNA-seq [27], and MAD-based selection is preferable to variance-based selection given its robustness to the outlier samples common in platelet transcriptomics [11]. That said, overlap persists across all three dimensionality reduction projections, indicating that cancer-induced TEP reprogramming includes substantial non-specific oncological signals shared across tumour types - a pattern consistent with prior TEP transcriptomic literature [11, 9].

### 4.3 Differential Expression and Biological Signatures

The 3,138 genes upregulated in healthy donors relative to the pooled cancer cohort is a striking result, and one that aligns well with evidence that cancer suppresses immune-cell RNA content in TEPs, depleting lymphocyte-derived transcripts such as CD3D, IL7R, and CCR7 [11]. This systematic immune depletion likely reflects tumour-derived immunosuppressive factors that alter platelet RNA uptake and splicing activity in the peripheral circulation. The small DEG sets for hepatobiliary cancer (17 genes) and CRC (32 genes) reflect limited statistical power from small group sizes and biological similarity to adjacent gastrointestinal and hepatic malignancies. Breast cancer yielded 207 significant DEGs yet showed the lowest per-class recall across nearly all classifiers. This discordance between univariate DEG count and multivariate classification performance underscores a key methodological point: univariate filtering does not reliably identify the joint feature combinations most discriminative in a multi-class setting [34].

The GO enrichment results were biologically plausible for four cancer types. Oxygen and gas transport enrichment in CRC-TEPs, driven by haemoglobin-related genes including HBG2 and FECH, likely reflects erythroid gene transcripts sequestered during tumour-associated vascular remodelling. NSCLC VEGF pathway enrichment is consistent with platelet-mediated sequestration of tumour-derived VEGF-A, supporting pro-angiogenic signalling [10]. For pancreatic cancer, actin cytoskeletal enrichment likely reflects cancer-induced platelet activation mediated by pancreatic tumour-derived tissue factor. The absence of significant enrichment for breast and hepatobiliary cancers is attributable to the small DEG sets and, in the case of breast cancer, to molecular heterogeneity that dilutes subtype-specific pathway signals when the disease is treated as a single class.

### 4.4 Machine Learning Performance and Statistical Reliability

Logistic Regression with multinomial SAGA optimization was the best overall classifier (test macro F1: 0.522; AUC: 0.869). While these values are moderate in absolute terms, both are substantially above the seven-class chance level (≈ 0.14) and demonstrate meaningful discrimination in a highly challenging multi-class liquid biopsy setting with limited training data. The superiority of regularised linear classification over more complex ensemble and deep learning models is consistent with the well-documented “p ≫ n” phenomenon: when the sample-to-feature ratio is small (n/p = 224/5,000 = 0.045), linear models with appropriate regularisation tend to generalise more robustly than high-capacity non-linear classifiers [34]. The class-specific AUC values ranged from 0.714 to 1.000 (macro-average: 0.869), indicating well-calibrated probabilistic output. Soft-probability-based decision rules may therefore be more appropriate for clinical deployment than hard argmax predictions. Bootstrap CI analysis [23, 24] confirmed that all model F1 and accuracy CIs are wide (typical width ≈ 0.25), reflecting the constraint of the small test partition (n = 56) rather than model instability. AUC CIs were narrower (width ≈ 0.12), confirming that AUC is the more reliably estimable metric in small-sample probabilistic settings. Most pair-wise model performance differences are not statistically resolvable at this test-set size: the sole exception is the Logistic Regression versus KNN comparison, where the AUC lower bound (0.804) does not overlap KNN’s upper bound (0.810). Relative model ranking should be treated as preliminary pending evaluation on a substantially larger independent cohort.

XGBoost achieved the highest precision (0.567) but showed a discrepancy between cross-validation and test accuracy (Δ ≈ 0.10), suggesting moderate overfitting to the training set [14]. LightGBM underperformed relative to XGBoost (F1: 0.364 vs. 0.469), possibly because its leaf-wise growth strategy overfits on small training partitions [15]. A consistent pattern across all classifiers was near-zero or very low breast cancer recall, with systematic misclassification towards pancreatic cancer - indicating genuine transcriptomic overlap between these classes in the TEP feature space that cannot be resolved without additional discriminative modalities. Hepatobiliary cancer recall was high across all models (0.33-1.00) but rests on only three test samples and must therefore be interpreted with caution.

### 4.5 Deep Learning vs. Classical Machine Learning

Despite its greater architectural capacity, the MLP performed below the top classical models in hard-label metrics (F1: 0.435 vs. 0.522 for Logistic Regression), though its AUC (0.866) was comparable to the top classical performers. The progressive training-validation divergence - training accuracy approaching ∼100% against validation accuracy plateauing at 65-70% - is consistent with deep networks memorising small training sets in the n ≪ p regime [33]. The regularisation strategies employed (Dropout 0.50/0.50/0.30, L2 weight decay λ = 10⁻⁴, and early stopping) partially mitigated but did not eliminate this behaviour [31, 32]. That the MLP’s AUC matched the top classical models suggests it acquired a useful probabilistic representation; translating this into reliable hard-label decisions would likely require dimensionality reduction at the input layer, variational autoencoder pre-training, or more aggressive cross-validated hyperparameter optimisation.

### 4.6 SHAP-Based Biological Interpretability

SHAP analysis of the Random Forest classifier identified both pan-cancer drivers and cancer-type-specific features [21]. IFITM3 (interferon-induced transmembrane protein 3) was the globally dominant gene and the leading discriminator for CRC and GBM. IFITM3 is an interferon-stimulated gene with established roles in innate antiviral immunity whose aberrant expression has been reported in multiple solid tumour types; its prominence in TEPs is consistent with platelet uptake of tumour-derived IFN-signalling intermediates in the circulation [9]. MDM4, a critical negative regulator of the p53 tumour suppressor pathway - featured prominently in the global top-10 and in the GBM-specific signature, in line with the frequent MDM4 amplification reported in glioblastoma. ATP5PD was the leading hepatobiliary discriminator, reflecting elevated mitochondrial oxidative phosphorylation in hepatobiliary cancer-educated platelets. C6orf62 ranked second globally and first for both NSCLC and pancreatic cancer; as a functionally under-characterised open reading frame whose importance is consistent across cancer types, it represents a high-priority candidate biomarker warranting prospective experimental validation.

### 4.7 Limitations and Future Directions

The sample size is modest relative to feature dimensionality, particularly for hepatobiliary cancer (n = 14), limiting the statistical reliability of class-specific conclusions. All analyses were conducted on the single publicly available TEP dataset (GSE68086) [11], the absence of external validation constrains the generalisability of the reported classifiers. The differential expression analysis used one-versus-rest Welch t-tests without covariate adjustment for patient age, sex, or clinical stage, each of which may independently influence the TEP transcriptome. Bootstrap CI analysis confirms that the small test set (n = 56) precludes statistically powered pair-wise model comparisons, underscoring the need for a substantially larger independent validation cohort.

Future work should prioritise:

1. Prospective external validation in independent TEP RNA-seq cohorts.
2. Integration of platelet proteomics, miRNA profiles, and clinical metadata as complementary multimodal features.
3. Application of attention-based or graph neural network architectures that can exploit gene-gene interaction structure.
4. Prospective clinical validation of top SHAP-identified biomarkers - particularly IFITM3, C6orf62, and ATP5PD - as platelet-based liquid biopsy targets.
5. Tumour-of-origin analysis to enable early cancer detection and localisation from a single blood draw.

## 5. Conclusion

TEP RNA-seq profiles from the GSE68086 dataset support automated multi-class cancer classification across six cancer types and healthy donors. Multinomial Logistic Regression with SAGA optimisation was the best-performing classifier, attaining a macro F1-score of 0.522 and macro-averaged ROC-AUC of 0.869 on the held-out test set - substantially above the seven-class chance level (≈ 0.14) - with class-specific AUC values ranging from 0.714 to 1.000. Bootstrap CI analysis (n = 2,000) [23, 24] confirmed the AUC advantage of Logistic Regression over KNN as the only statistically supported pair-wise distinction at the current test-set size, motivating validation in larger independent cohorts.

SHAP analysis identified IFITM3 as the globally dominant TEP biomarker, with cancer-type-specific drivers including ATP5PD (hepatobiliary cancer), C6orf62 (NSCLC and pancreatic cancer), VPS13C (healthy donors), and TMSB4Y (breast cancer). GO and KEGG enrichment analysis corroborated the biological plausibility of the identified differential expression signatures; the NSCLC enrichment for the KEGG “Non-Small Cell Lung Cancer” pathway directly validates the biological specificity of the computational pipeline. Taken together, these findings support the diagnostic potential of TEP transcriptomics as a minimally invasive, blood-based platform for pan-cancer screening, and highlight the importance of model interpretability in translating machine learning classifiers into clinically actionable liquid biopsy tools. Advances in dataset scale, multi-omics integration, and prospective clinical study design will be necessary to realise this potential in practice.

## 6. Data and Code Availability

The RNA-seq transcriptomic dataset analyzed in this study is publicly available through the NCBI Gene Expression Omnibus (GEO) under accession number GSE68086. The complete analysis pipeline, preprocessing scripts, machine learning workflows, differential expression analysis code, visualization scripts, and supplementary computational resources used in this study are openly available on GitHub at: ML-based Pan-Cancer Classification from TEP RNA-Seq GitHub Repository

## Acknowledgement

We thank the authors of the original GSE68086 study for making the tumor-educated platelet RNA-seq dataset publicly available.

---

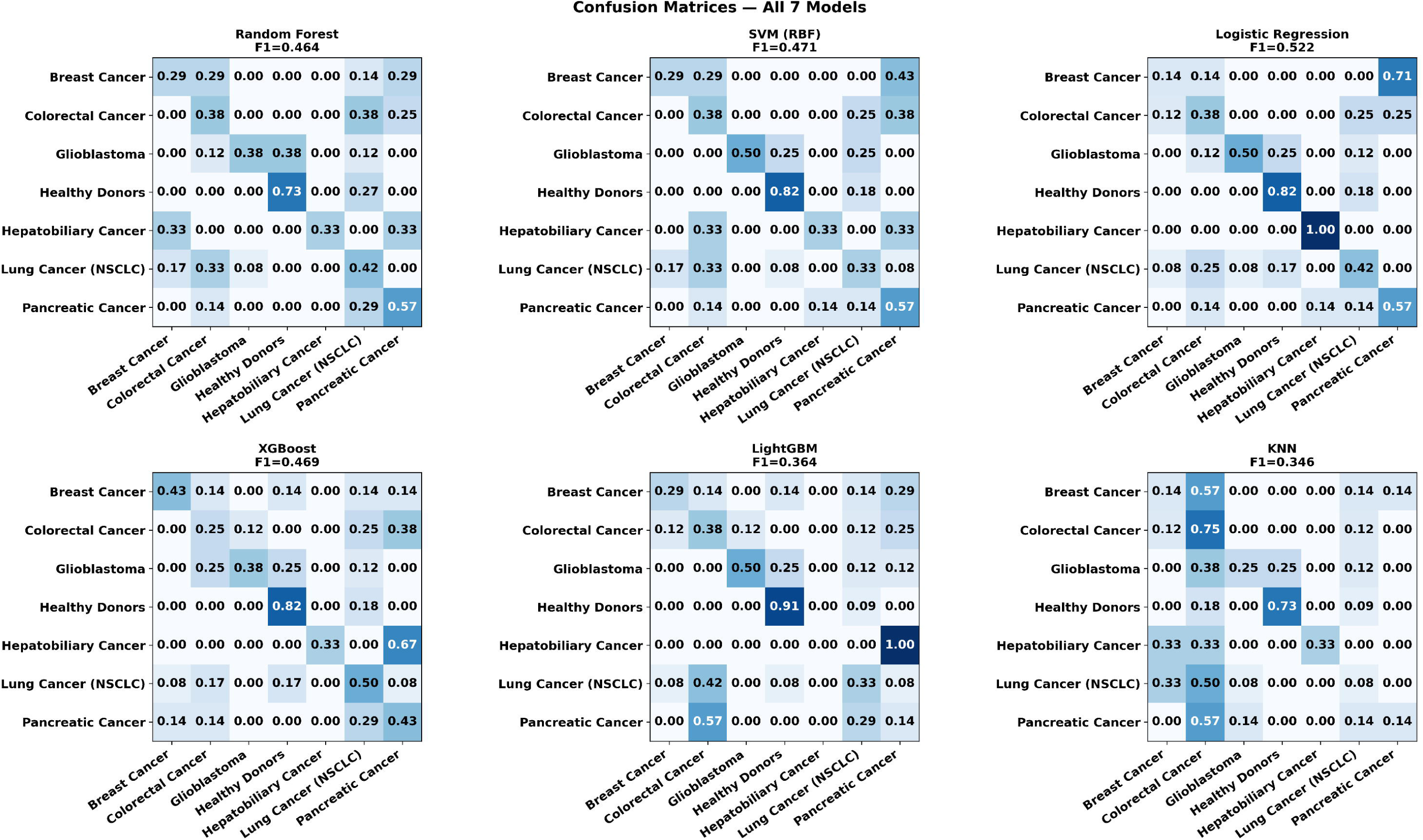

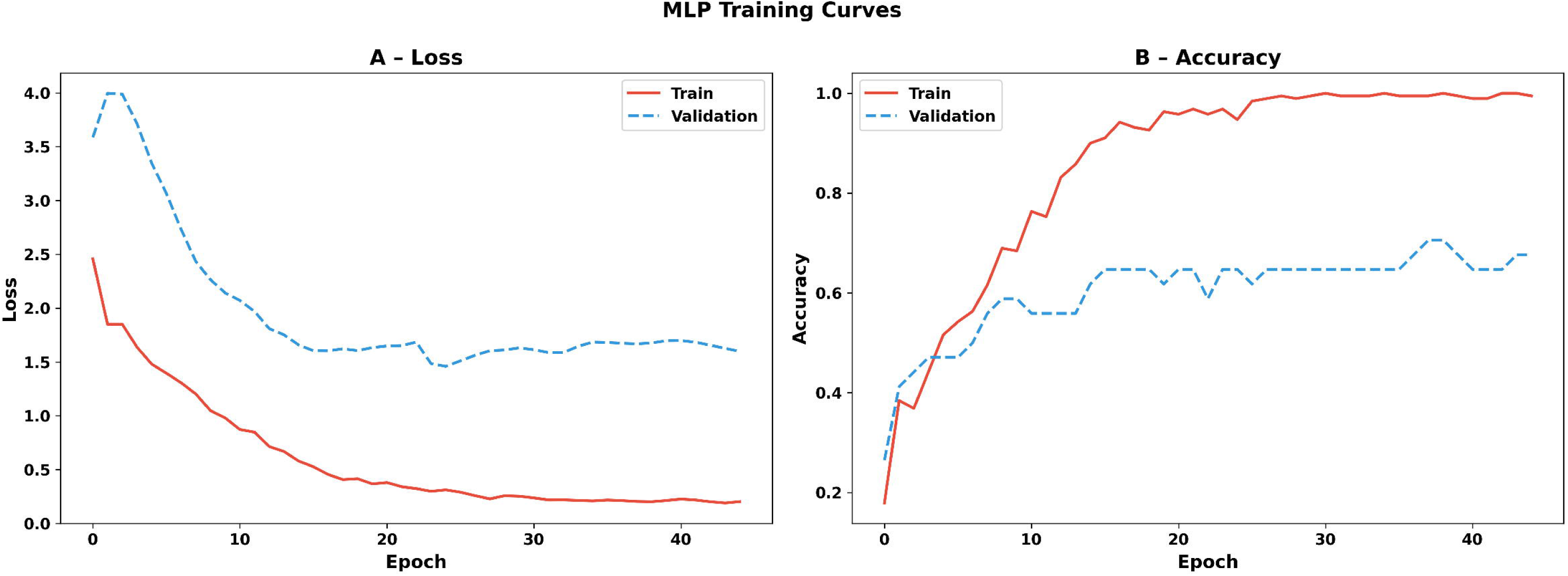

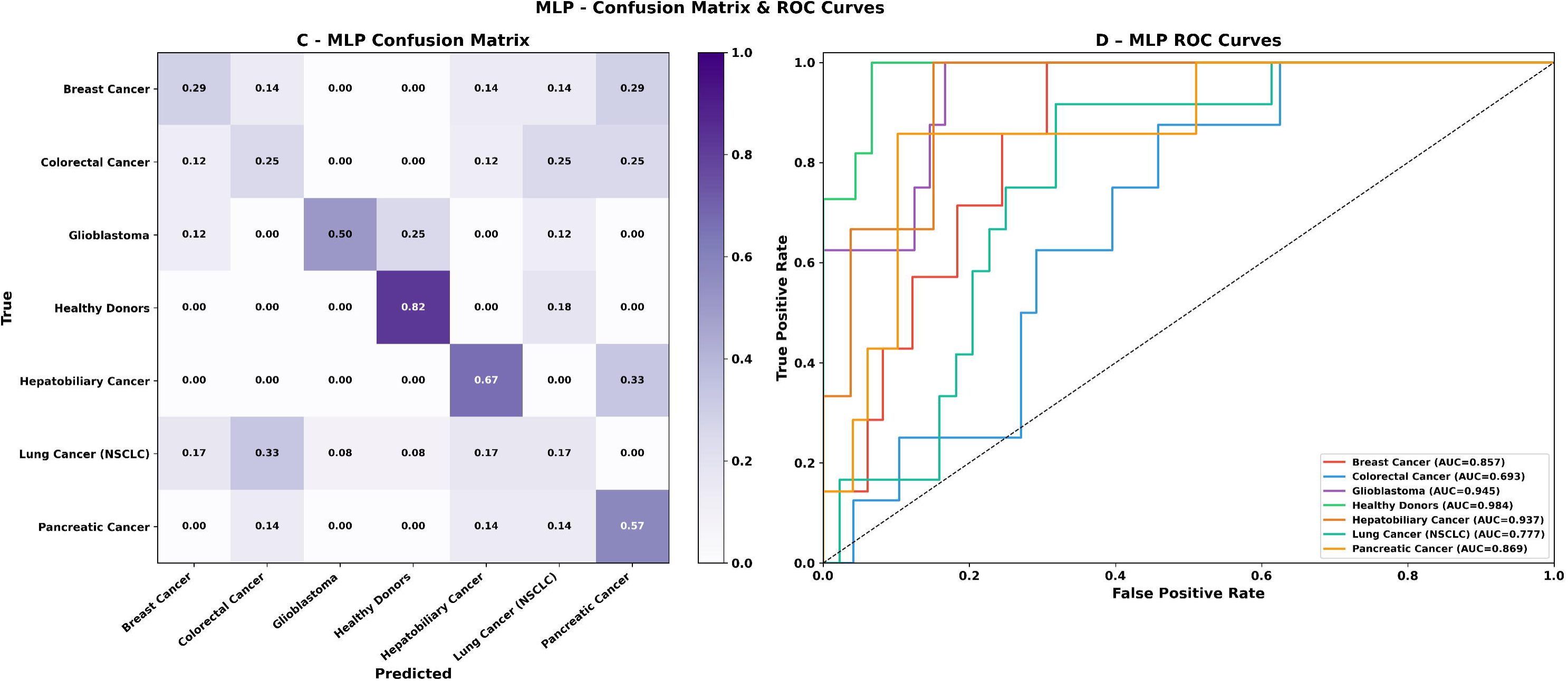

## References

[1] Bray, Freddie, et al. “Global Cancer Statistics 2022: GLOBOCAN Estimates of Incidence and Mortality Worldwide for 36 Cancers in 185 Countries.” CA: A Cancer Journal for Clinicians, vol. 74, no. 3, 2024, pp. 229–63. PubMed, 10.3322/caac.21834.

[2] Siegel, Rebecca L., et al. “Cancer Statistics, 2023.” CA: A Cancer Journal for Clinicians, vol. 73, no. 1, Jan. 2023, pp. 17–48. PubMed, 10.3322/caac.21763.

[3] Bettegowda, Chetan, et al. “Detection of Circulating Tumor DNA in Early- and Late-Stage Human Malignancies.” Science Translational Medicine, vol. 6, no. 224, Feb. 2014, p. 224ra24. PubMed, 10.1126/scitranslmed.3007094.

[4] Wan, Jonathan C. M., et al. “Liquid Biopsies Come of Age: Towards Implementation of Circulating Tumour DNA.” Nature Reviews. Cancer, vol. 17, no. 4, Apr. 2017, pp. 223–38. PubMed, 10.1038/nrc.2017.7.

[5] Gay, Laurie J., and Brunhilde Felding-Habermann. “Contribution of Platelets to Tumour Metastasis.” Nature Reviews. Cancer, vol. 11, no. 2, Feb. 2011, pp. 123–34. PubMed, 10.1038/nrc3004.

[6] Labelle, Myriam, et al. “Direct Signaling between Platelets and Cancer Cells Induces an Epithelial-Mesenchymal-like Transition and Promotes Metastasis.” Cancer Cell, vol. 20, no. 5, Nov. 2011, pp. 576–90. PubMed, 10.1016/j.ccr.2011.09.009.

[7] Nilsson, R. Jonas A., et al. “Blood Platelets Contain Tumor-Derived RNA Biomarkers.” Blood, vol. 118, no. 13, Sept. 2011, pp. 3680–83. PubMed, 10.1182/blood-2011-03-344408.

[8] Rowley, Jesse W., et al. “Genome-Wide RNA-Seq Analysis of Human and Mouse Platelet Transcriptomes.” Blood, vol. 118, no. 14, Oct. 2011, pp. e101–111. PubMed, 10.1182/blood-2011-03-339705.

[9] Khoo, Bee Luan, et al. “Liquid Biopsy and Therapeutic Response: Circulating Tumor Cell Cultures for Evaluation of Anticancer Treatment.” Science Advances, vol. 2, no. 7, July 2016, p. e1600274. DOI.org (Crossref), 10.1126/sciadv.1600274.

[10] Khoo, Bee Luan, et al. “Liquid Biopsy and Therapeutic Response: Circulating Tumor Cell Cultures for Evaluation of Anticancer Treatment.” Science Advances, vol. 2, no. 7, July 2016, p. e1600274. PubMed, 10.1126/sciadv.1600274.

[11] Best, Myron G., et al. “RNA-Seq of Tumor-Educated Platelets Enables Blood-Based Pan-Cancer, Multiclass, and Molecular Pathway Cancer Diagnostics.” Cancer Cell, vol. 28, no. 5, Nov. 2015, pp. 666–76. PubMed, 10.1016/j.ccell.2015.09.018.

[12] Barrett, Tanya, et al. “NCBI GEO: Archive for Functional Genomics Data Sets--Update.” Nucleic Acids Research, vol. 41, no. Database issue, Jan. 2013, pp. D991–995. PubMed, 10.1093/nar/gks1193.

[13] Breiman L. Random forests. Mach Learn. 2001;45(1):5–32.

[14] Chen, Tianqi, and Carlos Guestrin. “XGBoost: A Scalable Tree Boosting System.” Proceedings of the 22nd ACM SIGKDD International Conference on Knowledge Discovery and Data Mining [San Francisco California USA], 2016, pp. 785–94. DOI.org (Crossref), 10.1145/2939672.2939785.

[15] Ke G, Meng Q, Finley T, Wang T, Chen W, Ma W, et al. LightGBM: A highly efficient gradient boosting decision tree. In: Advances in Neural Information Processing Systems; 2017; Long Beach, CA, USA. Vol. 30. p. 3146–3154.

[16] Kourou, Konstantina, et al. “Machine Learning Applications in Cancer Prognosis and Prediction.” Computational and Structural Biotechnology Journal, vol. 13, 2015, pp. 8–17. PubMed, 10.1016/j.csbj.2014.11.005.

[17] LeCun, Yann, et al. “Deep Learning.” Nature, vol. 521, no. 7553, May 2015, pp. 436–44. DOI.org (Crossref), 10.1038/nature14539.

[18] Eraslan, Gökcen, et al. “Deep Learning: New Computational Modelling Techniques for Genomics.” Nature Reviews. Genetics, vol. 20, no. 7, July 2019, pp. 389–403. PubMed, 10.1038/s41576-019-0122-6.

[19] Hoadley, Katherine A., et al. “Cell-of-Origin Patterns Dominate the Molecular Classification of 10,000 Tumors from 33 Types of Cancer.” Cell, vol. 173, no. 2, Apr. 2018, pp. 291–304.e6. PubMed, 10.1016/j.cell.2018.03.022.

[20] Rudin, Cynthia. “Stop Explaining Black Box Machine Learning Models for High Stakes Decisions and Use Interpretable Models Instead.” Nature Machine Intelligence, vol. 1, no. 5, May 2019, pp. 206–15. PubMed, 10.1038/s42256-019-0048-x.

[21] Lundberg, Scott, and Su-In Lee. “A Unified Approach to Interpreting Model Predictions.” arXiv:1705.07874, arXiv, 25 Nov. 2017. arXiv.org, 10.48550/arXiv.1705.07874.

[22] Lundberg, Scott M., et al. “From Local Explanations to Global Understanding with Explainable AI for Trees.” Nature Machine Intelligence, vol. 2, no. 1, Jan. 2020, pp. 56–67. PubMed Central, 10.1038/s42256-019-0138-9.

[23] Efron, Bradley. “Bootstrap Methods: Another Look at the Jackknife.” Breakthroughs in Statistics: Methodology and Distribution, edited by Samuel Kotz and Norman L. Johnson, Springer, 1992, pp. 569–93. Springer Link, 10.1007/978-1-4612-4380-9_41.

[24] Efron, Bradley, and Robert J. Tibshirani. An Introduction to the Bootstrap. Springer US, 1993. DOI.org (Crossref), 10.1007/978-1-4899-4541-9.

[25] Benjamini, Y., et al. “Controlling the False Discovery Rate in Behavior Genetics Research.” Behavioural Brain Research, vol. 125, nos. 1-2, Nov. 2001, pp. 279–84. PubMed, 10.1016/s0166-4328(01)00297-2.

[26] Kuleshov, Maxim V., et al. “Enrichr: A Comprehensive Gene Set Enrichment Analysis Web Server 2016 Update.” Nucleic Acids Research, vol. 44, no. W1, July 2016, pp. W90–97. PubMed, 10.1093/nar/gkw377.

[27] Robinson, Mark D., et al. “edgeR: A Bioconductor Package for Differential Expression Analysis of Digital Gene Expression Data.” *Bioinformatics (Oxford*, England), vol. 26, no. 1, Jan. 2010, pp. 139–40. PubMed, 10.1093/bioinformatics/btp616.

[28] Welch, B. L. “The Generalisation of Student’s Problems When Several Different Population Variances Are Involved.” Biometrika, vol. 34, nos. 1-2, 1947, pp. 28–35. PubMed, 10.1093/biomet/34.1-2.28.

[29] Cortes C, Vapnik V. Support-vector networks. Mach Learn. 1995;20(3):273–297.

[30] Murphy, O. J. “Nearest Neighbor Pattern Classification Perceptrons.” Proceedings of the IEEE, vol. 78, no. 10, Oct. 1990, pp. 1595–98. IEEE Xplore, 10.1109/5.58344.

[31] Abadi, Martín, et al. “TensorFlow: A System for Large-Scale Machine Learning.” arXiv, 2016. DOI.org (Datacite), 10.48550/ARXIV.1605.08695.

[32] Ioffe, Sergey, and Christian Szegedy. “Batch Normalization: Accelerating Deep Network Training by Reducing Internal Covariate Shift.” arXiv:1502.03167, arXiv, 2 Mar. 2015. arXiv.org, 10.48550/arXiv.1502.03167.

[33] Ali, Shahwan Younis, and Hajar Maseeh. “Dropout: An Effective Approach to Prevent Neural Networks from Overfitting.” Asian Journal of Research in Computer Science, vol. 18, no. 2, Feb. 2025, pp. 163–85. DOI.org (Crossref), 10.9734/ajrcos/2025/v18i2569.

[34] Pedregosa, Fabian, et al. “Scikit-Learn: Machine Learning in Python.” Journal of Machine Learning Research, vol. 12, no. 85, 2011, pp. 2825–30. www.jmlr.org, http://jmlr.org/papers/v12/pedregosa11a.html.

[35] Fawcett, Tom. “An Introduction to ROC Analysis.” Pattern Recognition Letters, vol. 27, no. 8, June 2006, pp. 861–74. DOI.org (Crossref), 10.1016/j.patrec.2005.10.010.

[36] Pearson, Karl. “LIII. *On Lines and Planes of Closest Fit to Systems of Points in Space*.” *The London*, Edinburgh, and Dublin Philosophical Magazine and Journal of Science, vol. 2, no. 11, Nov. 1901, pp. 559–72. DOI.org (Crossref), 10.1080/14786440109462720.

[37] Hotelling, H. “Analysis of a Complex of Statistical Variables into Principal Components.” Journal of Educational Psychology, vol. 24, no. 6, Sept. 1933, pp. 417–41. DOI.org (Crossref), 10.1037/h0071325.

[38] McInnes, Leland, et al. “UMAP: Uniform Manifold Approximation and Projection.” Journal of Open Source Software, vol. 3, no. 29, Sept. 2018, p. 861. joss.theoj.org, 10.21105/joss.00861.

[39] Maaten, Laurens van der, and Geoffrey Hinton. “Visualizing Data Using T-SNE.” Journal of Machine Learning Research, vol. 9, no. 86, 2008, pp. 2579–605. www.jmlr.org, http://jmlr.org/papers/v9/vandermaaten08a.html.

[40] Kingma, Diederik P., and Jimmy Ba. “Adam: A Method for Stochastic Optimization.” arXiv.Org, 22 Dec. 2014, https://arxiv.org/abs/1412.6980v9.

